# Mitotic phosphorylation of ADAR1 regulates its centromeric localization and is required for faithful mitotic progression

**DOI:** 10.1101/2025.05.28.656747

**Authors:** Yuxi Yang, Mai Kubota, Kokone Hasegawa, Chao Zeng, Kazuko Nishikura, Michiaki Hamada, Masayuki Sakurai

## Abstract

Adenosine Deaminase Acting on RNA 1 p110 (ADAR1p110), the constitutively expressed nuclear isoform of the RNA-editing enzyme ADAR1, plays well-established roles in adenosine-to-inosine RNA editing, but its functions during mitosis remain poorly defined. Here, we identify ADAR1p110 as a chromatin-associated factor essential for mitotic chromosome segregation and cell viability. Depletion of ADAR1 caused metaphase arrest, DNA damage accumulation, and apoptosis. Co-immunoprecipitation experiments revealed that ADAR1p110 physically interacts with Structural Maintenance of Chromosome 3 (SMC3), a core component of the cohesin complex. Genome-wide DNA immunoprecipitation sequencing (DIP-seq) showed that ADAR1p110 selectively binds centromeric α-satellite DNA during mitosis, a finding validated by DNA immunoprecipitation quantitative polymerase chain reaction (DIP-qPCR). Complementary DNA-RNA immunoprecipitation sequencing (DRIP-seq) revealed that RNA:DNA hybrids (R-loops) are enriched in centromeric regions during mitosis and are further augmented by ADAR1 overexpression. We identified serine 614 (S614) as a key mitosis-specific phosphorylation site on ADAR1p110, and demonstrated that this post-translational modification is essential for its chromatin recruitment, stability, and mitotic function. Rescue experiments using phospho-mimetic mutants (S614D and 3×D) successfully restored mitotic progression in ADAR1-deficient cells, whereas non-phosphorylatable variants failed to do so. These results reveal that ADAR1p110 is phosphorylated in a cell cycle-dependent manner and functions at the intersection of post-transcriptional and post-translational regulation. By coordinating R-loop recognition at centromeres with chromosome cohesion, ADAR1p110 safeguards genome integrity during mitosis. Our findings uncover a previously uncharacterized mechanism through which a canonical RNA-editing enzyme contributes to chromosomal dynamics independent of its deaminase activity.

## Introduction

Adenosine-to-inosine (A-to-I) RNA editing, catalyzed by Adenosine Deaminase Acting on RNA (ADAR) enzymes, is a widespread post-transcriptional mechanism that alters RNA sequence, structure, and function (Nishikura 2010). Among the ADAR family, ADAR1 plays an essential role in mammalian development and immune homeostasis, and is best known for editing double-stranded RNAs (dsRNAs) to prevent activation of innate immune sensors (Pestal et al. 2015; Eisenberg and Levanon 2018; Samuel 2011). Two major ADAR1 isoforms are produced from alternative promoters: the interferon-inducible cytoplasmic isoform p150, and the constitutively expressed nuclear isoform p110 (Patterson and Samuel 1995; George and Samuel 1999). While ADAR1p150 has been well studied in the context of antiviral responses, the functions of nuclear-localized ADAR1p110 are only beginning to be elucidated.

Recent studies have expanded the known roles of ADAR1p110 beyond RNA editing. This isoform binds not only dsRNA, but also RNA:DNA hybrid structures (R-loops), implicating it in broader nuclear processes including chromatin regulation and genome surveillance (Shiromoto et al. 2021; Jimeno et al. 2021). R-loops, once considered transcriptional byproducts, are now recognized as functional elements that influence chromatin accessibility, transcriptional control, and genome stability (Niehrs and Luke 2020). In particular, centromeric R-loops have been linked to kinetochore assembly and sister chromatid cohesion, suggesting that they have a regulatory role during mitosis (Mishra et al. 2021).

Mitosis involves a dramatic reorganization of chromatin, during which transcription is globally silenced and chromosome segregation becomes highly coordinated (Prescott and Bender 1962; Martínez-Balbás et al. 1995; Palozola et al. 2017). Recent findings indicate that several RNA-binding proteins participate in mitotic chromatin architecture or function (Ito et al. 2020; Castello et al. 2012; Rajagopal et al. 2025; Remsburg et al. 2023), yet the involvement of ADAR1p110 in these processes remains poorly characterized. Of particular interest is the cohesin complex, which maintains sister chromatid cohesion and is regulated through its interaction with chromatin (Nasmyth and Haering 2009; Peters and Nishiyama 2012). Recent studies suggest that R-loops may stabilize cohesin at defined genomic regions, including centromeres, thereby contributing to mitotic fidelity (Racca et al. 2021; Kabeche et al. 2018). These observations raise the possibility that ADAR1p110 may influence mitotic chromosome dynamics via chromatin-associated interactions.

Post-translational modifications, particularly phosphorylation, are central to the regulation of mitotic events such as chromatin condensation, spindle assembly, and checkpoint control (Schmitz et al. 2020; Nigg 2001; Li et al. 2013). Mitotic kinases—including cyclin-dependent kinases (CDKs), Aurora kinases, and Polo-like kinases (PLKs)—orchestrate these processes by modulating the activity, localization, and stability of chromatin-bound regulatory proteins (Nigg 2001; Olsen et al. 2010; Lindqvist et al. 2009). Given that several nuclear RNA-binding proteins are known to undergo phosphorylation in a cell cycle-dependent manner (Parrott et al. 2005; Kim et al. 2008), we hypothesized that ADAR1p110 might be similarly regulated through phosphorylation, enabling its recruitment to mitotic chromatin and supporting chromosome cohesion.

To test this hypothesis, we investigated whether ADAR1p110 is regulated by mitosis-specific phosphorylation, and whether this modification influences its association with centromeric chromatin or its role in cohesion maintenance. This study reveals a phosphorylation-dependent mechanism via which ADAR1p110 contributes to mitotic progression, uncovering a previously unrecognized chromatin regulatory role for this RNA-binding enzyme.

## Results

### ADAR1 depletion induces mitotic arrest and DNA damage

In our previous study, we analyzed the impact of ADAR1 depletion on cell dynamics using live cell imaging (Shiromoto et al. 2021). Cells treated with control small interfering RNA (siRNA) progressed through the cell cycle normally, with mitosis completing within ∼1 h before transitioning into G1 phase. By contrast, cells treated with ADAR1 targeting siRNA exhibited altered cell cycle progression. Specifically, a delay or arrest in mitosis was frequently observed; instead of completing mitosis within 1 h, these cells remained arrested at metaphase for over 8 h, eventually undergoing apoptosis.

To further characterize this phenotype, we performed flow cytometric analysis of the cell cycle following ADAR1 knockdown **(Fig. 1A).** Cells were stained with DRAQ5 to assess DNA content and with an antibody against phospho-histone H3 (Ser10) to identify mitotic cells. At 72 h after siRNA transfection, the proportion of cells exhibiting 4N DNA content and positive staining for phospho-histone H3 was significantly higher in ADAR1 knockdown samples than in control samples. These findings were consistent with microscopy observations and indicate that suppression of ADAR1 leads to mitotic arrest.

**Figure 1.**
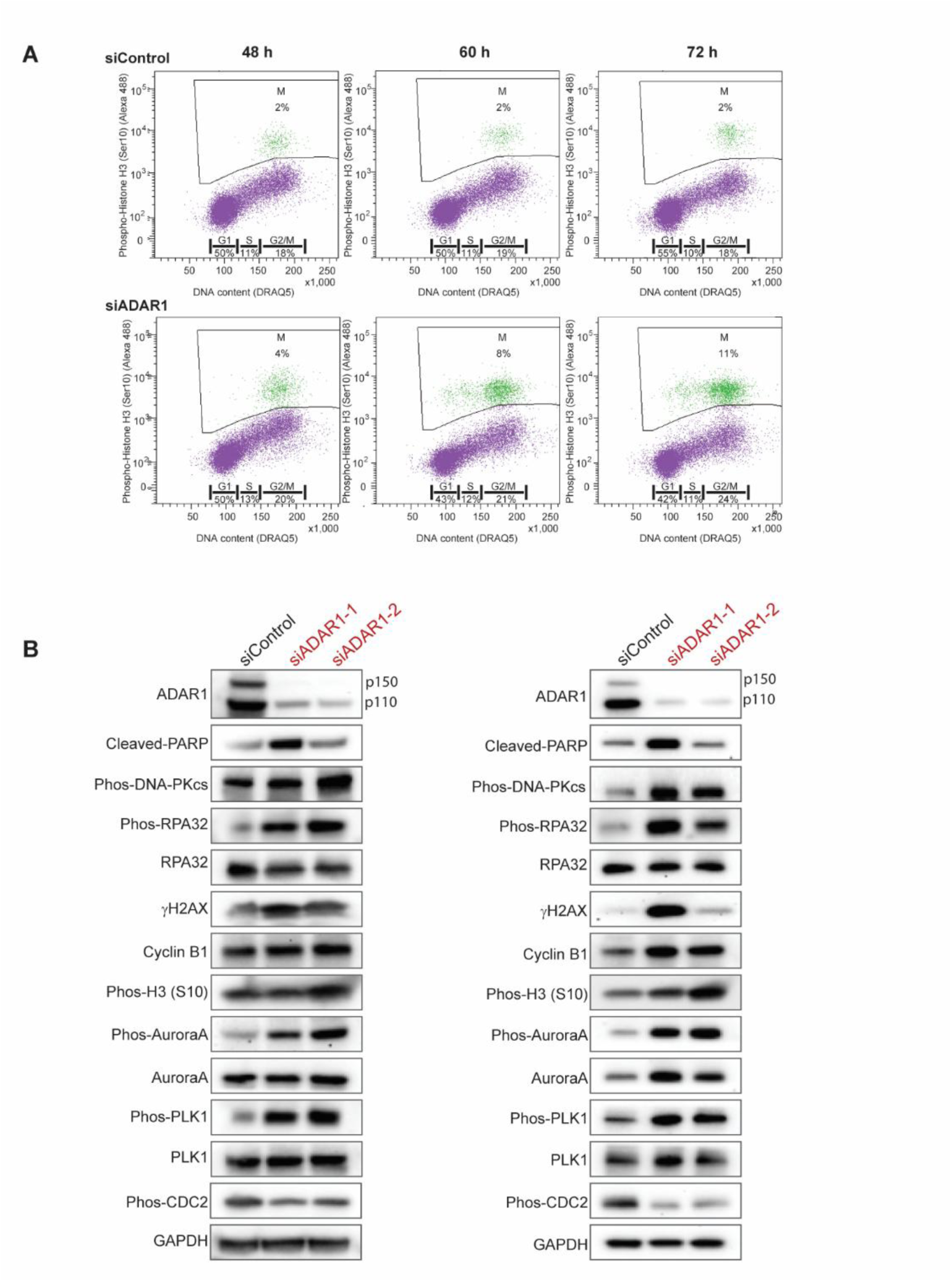
ADAR1 knockdown leads to DNA damage, apoptotic signaling, and mitotic marker accumulation. (A) Flow cytometry analysis showing DNA content (DRAQ5) on the x-axis and phospho-Histone H3 (Ser10) on the y-axis. HeLa cells were transfected with control siRNA (siControl, upper panels) or siRNA targeting ADAR1 (siADAR1, lower panels), and collected at 48, 60, and 72 h post-transfection (left to right). Percentages of cells in each cell cycle phase (G1, S, G2/M) and mitotic population (M, boxed area) are indicated. Green dots represent cells positive for phospho-Histone H3 (Ser10), indicating mitotic cells, while purple dots represent non-mitotic populations. (B) Western blotting analysis of mitotic and DNA damage markers following ADAR1 knockdown. HCT116 (left) and HeLa (right) cells were transfected with control siRNA (siControl) or two independent siRNAs targeting ADAR1 (siADAR1-1 and siADAR1-2). Protein lysates were collected and subjected to immunoblotting for ADAR1, cleaved PARP, phospho-DNA-PKcs, phospho-RPA32, RPA32, γH2AX, Cyclin B1, phospho-histone H3 (Ser10), phospho-PLK1 (T210), PLK1, phospho-Aurora A (T288), Aurora A, phospho-CDC2 (T15), and glyceraldehyde-3-phosphate dehydrogenase (GAPDH; loading control). Markers were selected to assess cell cycle status during mitosis and the presence of DNA damage.

To investigate the effects of ADAR1 depletion on mitotic progression and cellular stress responses, we performed western blotting analysis of HeLa and HCT116 cells following siRNA-mediated knockdown of ADAR1 **(Fig. 1B)**. In both cell lines, cleaved poly (ADP-ribose) polymerase (PARP) was increased in ADAR1-depleted cells, indicating enhanced cell death. Furthermore, levels of DNA damage markers including phosphorylated DNA-dependent protein kinase catalytic subunit (DNA-PKcs), phosphorylated replication protein A 32 kDa subunit (RPA32), and histone family member X (γH2AX) were elevated, suggesting the accumulation of DNA damage. Mitotic arrest was supported by observing upregulation of mitotic markers such as Cyclin B1, phospho-PLK1 (T210), phospho-Aurora A (T288), and phospho-histone H3 (Ser10). Additionally, dephosphorylation of CDC2 at T15 has been reported to correlate with mitotic progression, and its reduction upon ADAR1 knockdown may reflect a mitotic arrest phenotype.

Together, these findings suggest that ADAR1 knockdown induces mitotic arrest accompanied by DNA damage and apoptosis, likely via a mechanism involving impaired mitotic progression and checkpoint activation.

### ADAR1p110 associates with mitotic kinases and cohesin components during mitosis

To identify proteins that interact with ADAR1p110 during mitosis, we performed co-immunoprecipitation (co-IP) experiments using a Flag-tagged ADAR1p110 construct **(Fig. 2A)**. HeLa-Tet-On cells stably expressing 1×Flag-ADAR1p110 were induced with doxycycline to increase its exogenous expression. Simultaneously, siRNA targeting the 3′-untranslated region (UTR) of endogenous ADAR1 was introduced to suppress endogenous ADAR1 expression, allowing specific detection of exogenous Flag-ADAR1p110. Cells were harvested under asynchronous (Async) or mitotically synchronized (Msync) conditions, and IP was performed using anti-Flag antibody. Western blotting analysis revealed that several mitotic regulators, including phosphorylated Aurora kinases (A, B, and C), PLK1, but not CDC25C, were co-immunoprecipitated with Flag-ADAR1p110 under mitotic conditions. Importantly, Structural Maintenance of Chromosome 3 (SMC3) and SMC2—core components of the cohesin complex— were also enriched in the ADAR1p110 IP fractions under mitotic synchronization. Given the roles of SMC2/3 in sister chromatid cohesion and centromeric architecture (Peters and Nishiyama 2012; Hirano 2012), their interaction with ADAR1p110 suggests that ADAR1 may associate with the structural framework of mitotic chromosomes. Collectively, these findings indicate that ADAR1p110 forms mitosis-specific complexes.

**Figure 2.**
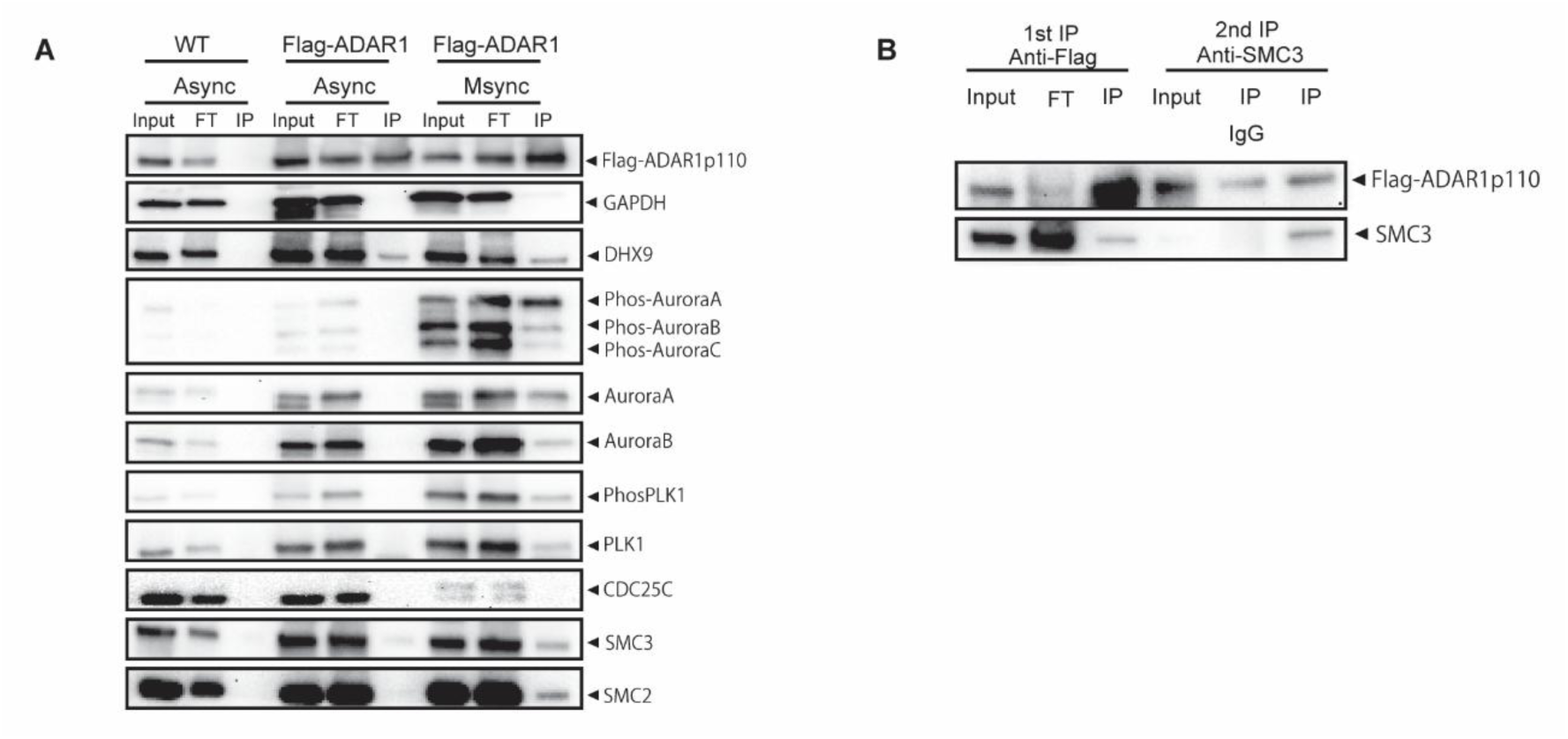
ADAR1p110 interacts with mitotic regulators and cohesin components during mitosis. (A) Identification of mitotic regulators interacting with ADAR1 by co-immunoprecipitation (co-IP). HeLa cells expressing Flag-tagged ADAR1p110 or wild-type (WT) controls were harvested under asynchronous (Async) or mitotically synchronized (Msync) conditions. Cell lysates were subjected to anti-Flag IP, and both input, flow-through (FT), and IP fractions were analyzed by western blotting. Immunoblots were probed for candidate interacting proteins, including DExD/H-box helicase (DHX9), Aurora kinases (Aurora A, B, C), PLK1, CDC25C, SMC3, SMC2, and phospho-specific forms of Aurora kinases and PLK1. GAPDH served as a negative control for nonspecific binding. (B) Lysates from Flag-ADAR1p110-expressing cells were subjected to IP using an anti-Flag antibody, and eluted protein complexes were re-immunoprecipitated using either anti-SMC3 or control IgG antibodies.

To further explore the mechanism via which ADAR1 functions during mitosis, we examined its potential interacting partners. Mass spectrometry analysis of ADAR1p110 immunoprecipitates from synchronized mitotic cells revealed several candidate proteins associated with chromosomal architecture, among which SMC3 was notably enriched. Given the critical role of cohesin in sister chromatid cohesion and chromosome segregation, we focused on the potential interaction between ADAR1 and SMC3. Immunofluorescence staining of endogenous ADAR1 and SMC3 revealed their dynamic co-localization during mitosis **(Supplemental Fig. 1)**. In prophase through anaphase, ADAR1p110 and SMC3 exhibited overlapping signals on chromosomes, suggesting that ADAR1 may associate with cohesin-enriched chromosomal regions.

To validate the physical interaction between ADAR1p110 and SMC3 observed in mitotically synchronized cells, we performed a sequential co-IP assay **(Fig. 2B)**. In the first step, lysates from Flag-ADAR1p110-expressing cells were immunoprecipitated using an anti-Flag antibody to isolate ADAR1-containing complexes. Eluates from the first IP were then subjected to a second round of IP using either anti-SMC3 or isotype control immunoglobulin G (IgG) antibodies. Western blotting analysis of the second IP showed that both Flag-ADAR1p110 and SMC3 were co-precipitated specifically in the anti-SMC3 IP fraction, while no signal was detected in the IgG control. These results confirmed that ADAR1p110 and SMC3 exist in the same protein complex, and that their interaction is specific rather than incidental.

These results support a model in which ADAR1p110 physically associates with the cohesin component SMC3 during mitosis. Given the known roles of cohesin in maintaining chromosomal architecture, these findings suggest that ADAR1 may contribute to chromosome dynamics through a cohesin-associated mechanism.

### ADAR1p110 localization to the mitotic spindle is dependent on its RNA-binding domain

To determine the subcellular localization of ADAR1p110 during mitosis, we performed fluorescence imaging in HeLa cells co-expressing mCherry-tagged wild-type (WT) ADAR1p110 and green fluorescent protein (GFP)-tagged α-tubulin **(Fig. 3A)**. DNA was counterstained with Hoechst 33342. Throughout metaphase, early anaphase, and late anaphase, WT ADAR1p110 was consistently observed along the mitotic spindle, suggesting a spatial association with spindle microtubules during cell division.

**Figure 3.**
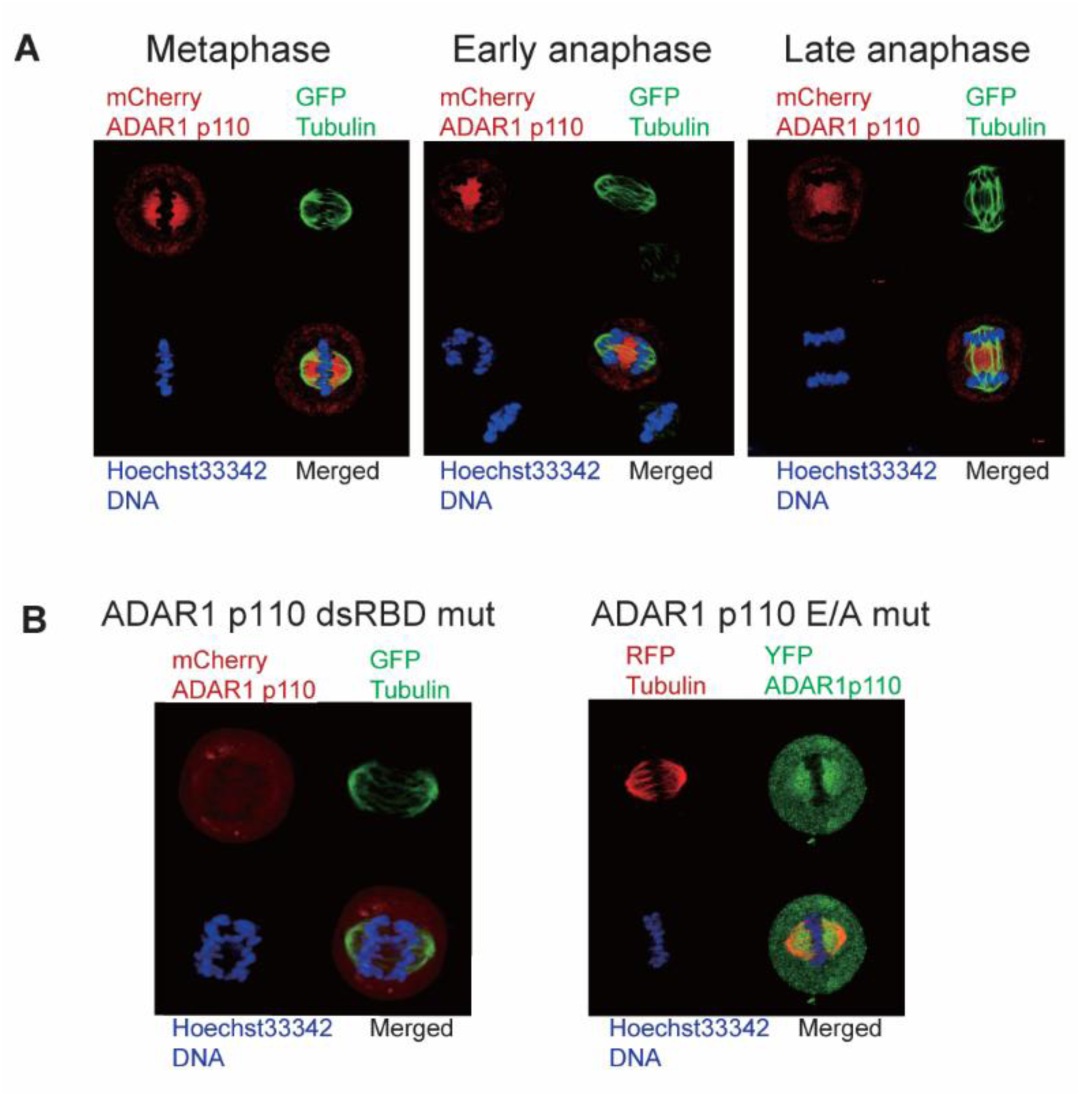
The RNA-binding domain is required for ADAR1p110 localization to the mitotic spindle. (A) Subcellular localization of ADAR1p110 during mitosis. HeLa cells were co-transfected with mCherry-tagged ADAR1p110 (red) and GFP-tagged α-tubulin (green) and imaged at distinct stages of mitosis (metaphase, early anaphase, and late anaphase). DNA was visualized by Hoechst 33342 staining (blue). Merged images demonstrate co-localization of ADAR1p110 with mitotic spindle microtubules. (B) Localization of mutant ADAR1p110 proteins during mitosis. HeLa cells expressing dsRNA-binding domain mutant (dsRBD mut; mCherry-ADAR1p110, red) or editing-inactive mutant (E/A mut; YFP-ADAR1p110, green) together with GFP- or red fluorescent protein (RFP)-tagged α-tubulin, respectively, were examined. DNA was stained with Hoechst 33342 (blue). The ADAR1p110 E/A mutant retained spindle localization, whereas the dsRBD mutant did not.

To investigate the domain requirements for this localization, we analyzed two functional mutants of ADAR1p110: a double-stranded RNA-binding domain (dsRBD) mutant defective in RNA binding, and an editing-inactive (E/A) mutant **(Fig. 3B)**. The E/A mutant retained spindle localization similar to WT protein, indicating that RNA editing activity is not essential for mitotic targeting. By contrast, the dsRBD mutant failed to associate with the spindle and remained localized to the nucleus.

These results indicate that ADAR1p110 localization to the mitotic spindle is mediated by its RBD, rather than its catalytic deaminase function. This supports a model in which ADAR1p110 is recruited to mitotic structures via RNA-mediated interactions rather than through enzymatic substrate recognition.

### ADAR1p110 preferentially associates with centromeric DNA during mitosis

ADAR1p110 is known to bind not only to dsRNA but also to RNA:DNA hybrid molecules, and to retain editing activity on such substrates (Jimeno et al. 2021; Shiromoto et al. 2021). Given that multiple mitotic regulators identified as ADAR1-interacting partners are known to associate with chromosomal DNA (Carmena et al. 2012; Kang et al. 2006), we hypothesized that ADAR1p110 may directly bind chromosomal DNA during mitosis. To explore this possibility, we performed DNA immunoprecipitation followed by next-generation sequencing (DNA-IP-seq, DIP-seq) targeting Flag-ADAR1p110 **(Fig. 4A)**. Cells were either left asynchronous or synchronized in mitosis prior to anti-Flag IP and DNA purification. Significant ADAR1-bound regions (*q* < 0.001) were mapped onto a telomere-to-telomere (T2T)-CHM13 genome karyogram. Under Async, 393 peaks were identified, whereas Msync cells exhibited 1,470 peaks, indicating a substantial increase in DNA binding during mitosis. Strikingly, ADAR1p110-binding sites in mitotic cells were found to be highly enriched in centromeric regions across multiple chromosomes. These results suggest that ADAR1p110 directly associates with specific genomic loci during mitosis, potentially linking its mitotic functions to centromeric chromatin. A genome-wide distribution of relaxed threshold peaks is shown in **Supplemental Figure 2**.

**Figure 4.**
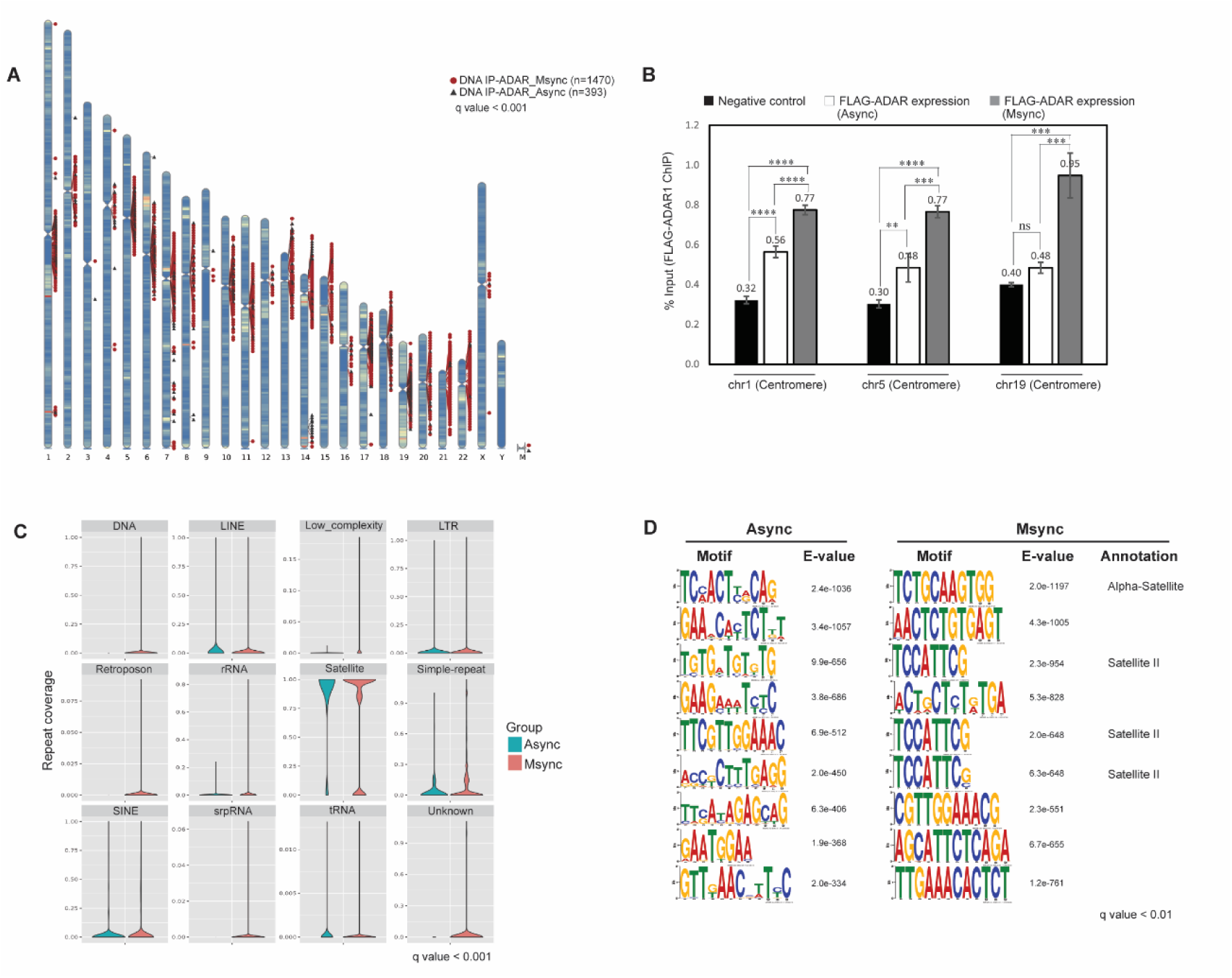
Genome-wide distribution of ADAR1p110-binding sites in asynchronous and mitotic cells. (A) DIP-seq was performed using HeLa cells expressing Flag-tagged ADAR1p110 under Async or Msync conditions. Significant ADAR1-bound DNA regions (*q* < 0.001) were mapped to the T2T-CHM13 reference genome and visualized on a karyogram. Red circles indicate ADAR1 peaks identified in mitotic cells (n = 1,470), and black triangles indicate peaks in asynchronous cells (n = 393). (B) DIP was performed using anti-Flag antibody in HeLa-Tet-On cells inducibly expressing Flag-tagged ADAR1p110, either under Async or Msync conditions. Quantitative PCR (qPCR) was carried out with primers targeting the centromeric regions of chromosomes 1, 5, and 19. Bars indicate the percentage of input DNA recovered in each IP: black bars represent negative control cells lacking Flag-ADAR1 expression, white bars represent asynchronous Flag-ADAR1-expressing cells, and gray bars represent mitotically synchronized Flag-ADAR1-expressing cells. Error bars show standard deviation (n = 3). Statistical analysis was performed using one-way analysis of variance (ANOVA) followed by Tukey’s post hoc test for multiple comparisons. *p* < 0.01 (**), *p* < 0.001 (***), *p* < 0.0001 (****). Results are mean ± standard error of the mean (SEM) from three independent experiments. (C) Violin plots showing the distribution of DIP-seq peak coverage across genomic repeat categories under Async (blue) and Msync (red) conditions. Repeat annotation was based on genome-wide RepeatMasker classification. ADAR1p110-bound peaks in mitotic cells showed significant enrichment in satellite repeats, particularly α-satellite and other centromere-associated elements. (D) *De novo* motif enrichment analysis was performed on DIP-seq peaks from Async (left) and Msync (right) cells. The top-ranked sequence motifs (based on E-value and site frequency) are shown as sequence logos. In mitotic cells, the dominant motifs corresponded to α-satellite and Satellite II repeats, while motifs from asynchronous cells were more heterogeneous and lacked centromeric annotation.

To validate the centromeric enrichment observed in DIP-seq, we conducted DNA IP followed by quantitative PCR (DIP-qPCR) using primers specific to the centromeric regions of chromosomes 1, 5, and 19—regions previously identified as ADAR1p110-binding sites **(Fig. 4B)**. DIP was performed using anti-Flag antibody in HeLa-Tet-On cells with inducible expression of 1×Flag-ADAR1p110 under either Async or Msync conditions. As a negative control, WT HeLa cells lacking Flag-ADAR1 expression were also subjected to DIP-qPCR. At all tested centromeric loci, qPCR revealed significant enrichment of ADAR1p110-bound DNA relative to the negative control. Furthermore, the level of enrichment was consistently higher in mitotically synchronized cells than asynchronous cells. This observation mirrors the global increase in centromeric association seen in the DIP-seq data and provides orthogonal validation that ADAR1p110 preferentially associates with centromeric DNA during mitosis.

To determine the sequence features of ADAR1p110-bound regions, we examined for the enrichment of repetitive elements within DIP-seq peaks **(Fig. 4C)**. Violin plot analysis revealed that satellite repeats were the most prominently enriched class in mitotic cells, with significantly higher coverage than in asynchronous cells. Among these, α-satellite (ALR-Alpha) sequences exhibited the strongest enrichment, followed by Satellite II elements. Notably, β-satellite (BSR) elements were detected only in mitotic samples, further supporting mitosis-specific binding of ADAR1p110.

To explore whether this enrichment reflects sequence specificity, we conducted *de novo* motif discovery on the ADAR1-bound regions **(Fig. 4D)**. In mitotically synchronized cells, the top motifs showed high similarity to consensus α-satellite and Satellite II sequences—known components of human centromeric DNA. By contrast, motifs enriched in asynchronous cells were more variable and lacked centromeric signatures.

These findings indicate that ADAR1p110 preferentially binds centromeric α-satellite DNA during mitosis, likely contributing to mitotic chromatin organization or stability. The specificity of this interaction suggests a potential regulatory role for ADAR1 in centromeric function during cell division.

### ADAR1p110 recruitment correlates with R-loop accumulation at centromeres during mitosis

To investigate the interplay between ADAR1p110 accumulation and R-loop formation at centromeres, we performed DNA-RNA immunoprecipitation sequencing (DRIP-seq) in Async or Msync cells, with and without ADAR1p110 overexpression. DRIP signals were consistently detectable in centromeric regions under all conditions, but notably increased in synchronized mitotic cells and further in those overexpressing ADAR1p110. These results suggest that R-loop formation at centromeres is both cell cycle-regulated and sensitive to ADAR1 expression levels.

To compare R-loop accumulation with ADAR1p110 chromatin association, we plotted the number of centromeric R-loop peaks (x-axis) against the normalized read count within centromeric regions (y-axis) for each condition **(Fig. 5A)**. ADAR1p110 DIP data (black squares) showed a mitosis-dependent increase in both the number and intensity of centromeric binding events. In parallel, DRIP-seq data (black circles) revealed increased centromeric R-loop formation under mitotic conditions and upon ADAR1 overexpression, suggesting a potential correlation between ADAR1p110 recruitment and R-loop accumulation at centromeres. To gain locus-specific insight, genome browser visualization of the chromosome 5 centromeric region revealed clear ADAR1p110 enrichment at α-satellite-like sequences, accompanied by increased R-loop signals in synchronized cells, especially upon ADAR1p110 overexpression **(Fig. 5B)**. This co-occurrence supports the idea that ADAR1p110 preferentially associates with R-loop-rich centromeric loci during mitosis.

**Figure 5.**
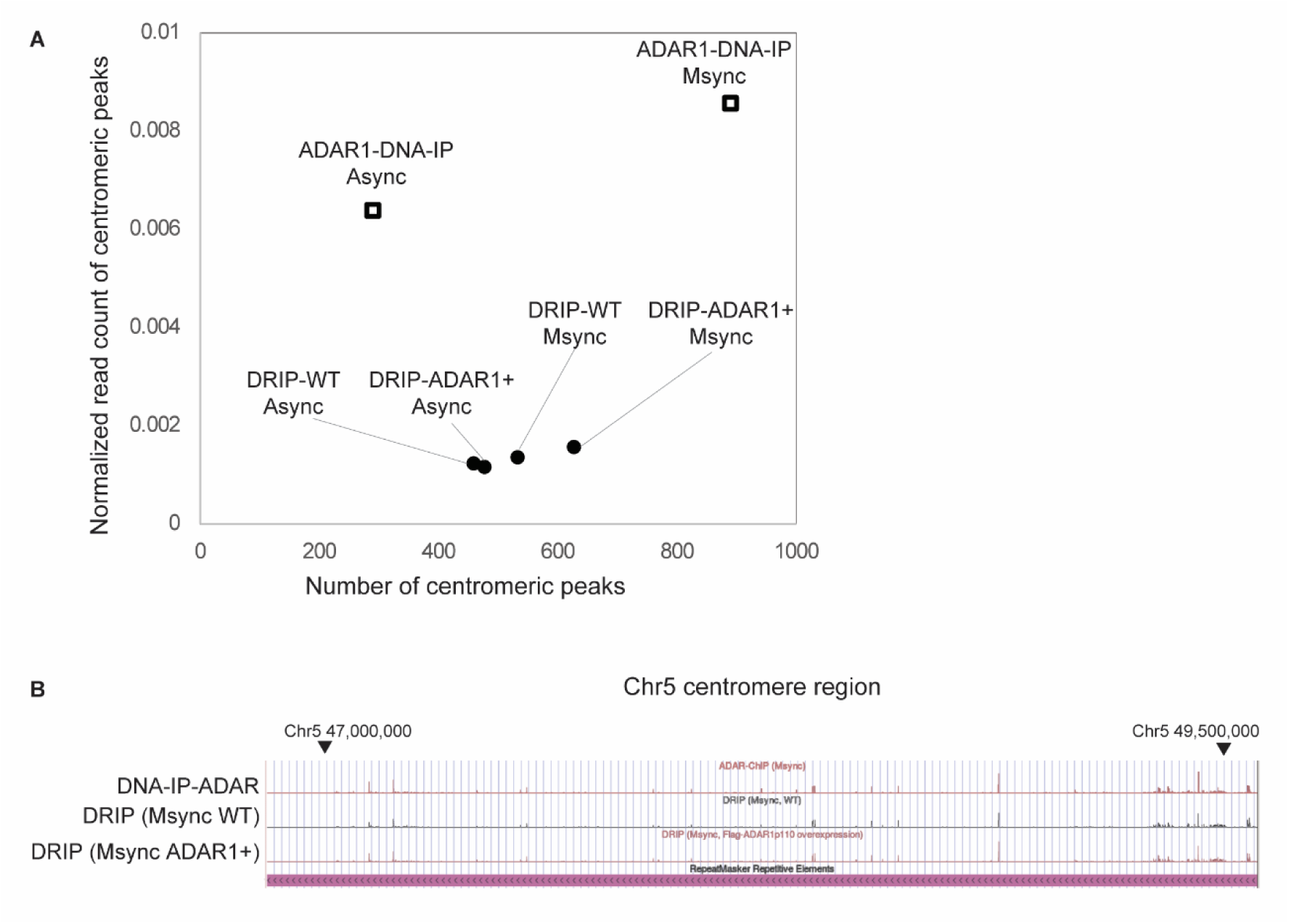
Comparative analysis of ADAR1p110 DNA binding and R-loop formation at centromeric regions. (A) Scatter plot comparing DIP (black squares) and DRIP-seq (black circles) data across Async and Msync conditions, with or without ADAR1p110 overexpression. The x-axis represents the proportion of the centromeric region covered by called peaks (peak coverage), and the y-axis shows normalized read counts within centromeres. ADAR1p110-binding (DIP) and R-loop signals (DRIP) were assessed to evaluate their respective enrichment patterns and potential co-localization dynamics under mitosis. (B) ADAR1p110 binds to R-loop-rich centromeric regions on chromosome 5 during mitosis. The genome browser view shows DIP and DRIP sequencing signals across the centromeric region of chromosome 5 (Chr5; 47,000,000−49,500,000). Top track: DIP-seq using anti-ADAR1 antibody in mitotically synchronized cells, showing ADAR1p110 enrichment at α-satellite-like sequences. Middle track: DRIP-seq signal from synchronized WT cells, reflecting endogenous R-loop distribution. Bottom track: DRIP-seq from synchronized cells overexpressing Flag-ADAR1p110, showing enhanced R-loop formation in the same region. The RepeatMasker track (bottom) indicates repetitive elements characteristic of the centromeric region.

Together, these observations suggest that centromeric R-loops are dynamically regulated during mitosis, and further augmented by ADAR1p110. The strong correlation between R-loop accumulation and ADAR1p110-binding supports a model in which ADAR1 may recognize or stabilize R-loop structures at centromeres to coordinate chromatin behavior during mitotic progression.

### Phosphorylation at serine 614 is required for ADAR1p110 to suppress mitotic accumulation

To determine whether ADAR1p110 is subject to phosphorylation in a cell cycle-regulated manner, we performed Phos-tag sodium dodecyl sulfate-polyacrylamide gel electrophoresis (SDS-PAGE) and western blotting on lysates from HeLa cells collected under asynchronous, M phase-arrested (nocodazole-treated), or S phase-arrested (thymidine-treated) conditions **(Fig. 6A)**. To verify the phosphorylation dependency of the observed mobility shifts, each sample was analyzed with or without λ-phosphatase treatment. In the Phos-tag gel (top panel), ADAR1p110 displayed a pronounced mobility shift in mitotically arrested cells, characterized by multiple slower-migrating bands. This shift was absent or minimal in asynchronous and S phase samples. Notably, λ-phosphatase treatment abolished the mobility shift across all conditions, confirming that the observed band pattern resulted from phosphorylation. These findings provide strong biochemical evidence that ADAR1p110 is preferentially phosphorylated during mitosis, and support the hypothesis that its phosphorylation is a mitosis-specific regulatory mechanism.

**Figure 6.**
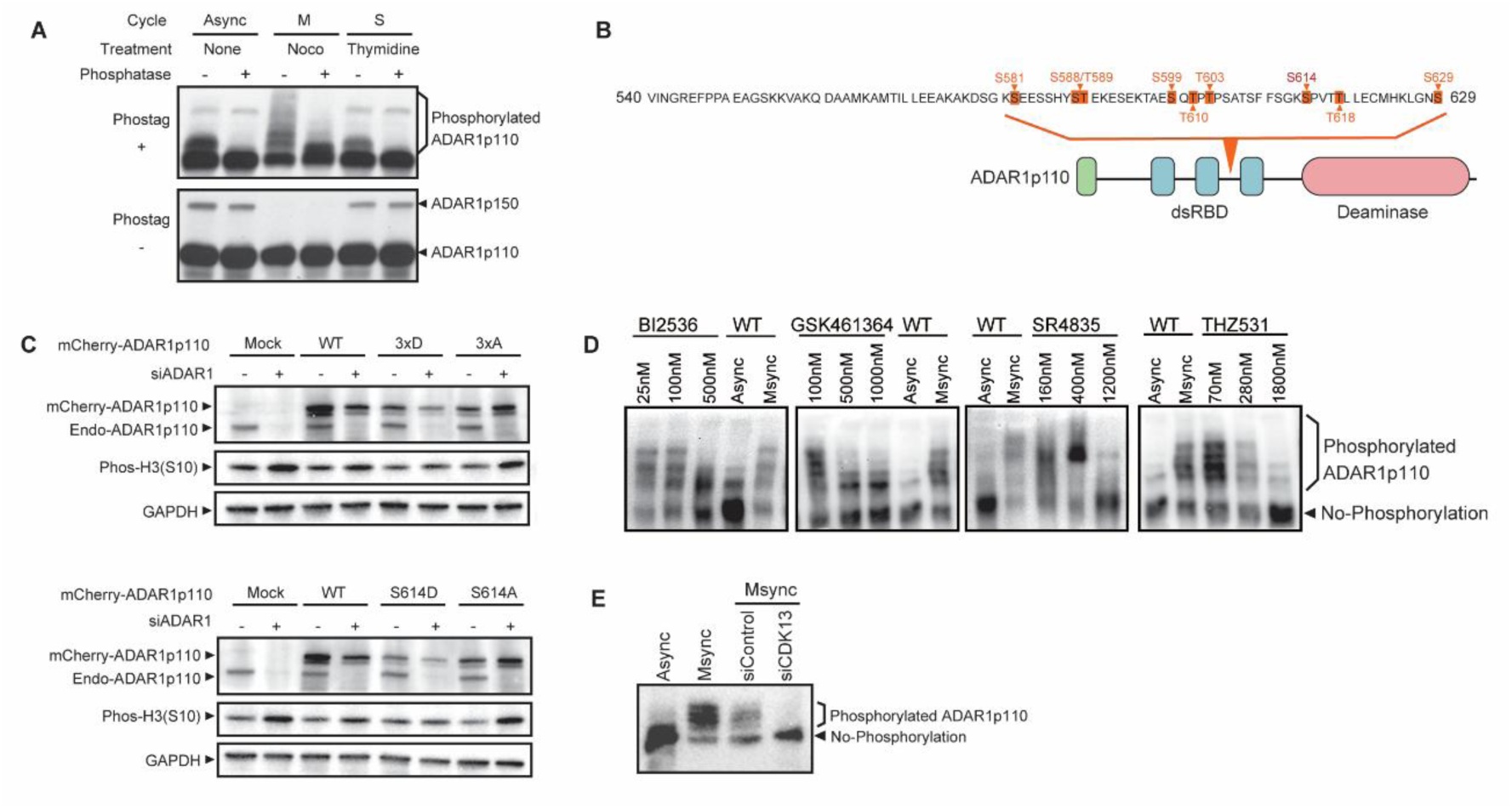
ADAR1p110 undergoes phosphorylation in a mitosis-specific manner. (A) HeLa cells were collected under asynchronous (Async), mitotically arrested (M; nocodazole-treated), or S phase-arrested (S; thymidine-treated) conditions. Cell lysates were analyzed by SDS-PAGE with (+) or without (–) Phos-tag acrylamide to detect phosphorylated ADAR1p110. λ-Phosphatase treatment was used to confirm phosphorylation dependency. In Phos-tag gels (top panel), ADAR1p110 exhibited a mobility shift that was strongly enhanced in mitotic samples, appearing as multiple slower-migrating bands. This shift was abolished by phosphatase treatment, indicating that the observed shift is phosphorylation-dependent. Conventional SDS-PAGE (bottom panel) was performed to assess total ADAR1p110 and ADAR1p150 protein levels as loading controls. (B) Mass spectrometry-based phosphopeptide mapping was performed on 3×Flag-tagged ADAR1p110 purified from 293T cells under mitotically synchronized conditions. The amino acid sequence starting from residue 514 is shown. Orange marks indicate phosphorylation sites. Below the sequence, a schematic representation of ADAR1p110 is provided, including the Z-DNA binding domain (green), the dsRBDs (blue), and the deaminase domain (red). (C) HeLa cells were transfected with siRNA targeting the 3′-untranslated region (3′-UTR) of ADAR1, followed by transfection with mCherry-tagged ADAR1p110 constructs. The constructs included WT, phospho-mimetic mutants (3×D and S614D), and phospho-deficient mutants (3×A and S614A). Cells were harvested 48 h after transfection, and total lysates were analyzed by western blotting using antibodies against mCherry (exogenous ADAR1p110), endogenous ADAR1p110, phospho-histone H3 (Ser10), and GAPDH. Phospho-histone H3 (S10) band intensity was used as a readout for mitotic accumulation under each condition. (D) HeLa cells were treated with kinase inhibitors under Async or Msync conditions. Cells were exposed to PLK1 inhibitors BI2536 and GSK461364, and CDK12/13 inhibitors SR-4835 and THZ531, across indicated concentrations. Whole-cell lysates were analyzed by Phos-tag SDS-PAGE followed by immunoblotting to detect phosphorylated ADAR1p110. Phosphorylated forms were visualized as slower-migrating bands. A decrease or disappearance of these bands indicates a loss of phosphorylation upon kinase inhibition. (E) HeLa cells were synchronized in Msync using nocodazole and transfected with either control siRNA (siNC1) or CDK13-targeting siRNA (siCDK13). Asynchronous cells were included for reference. Whole-cell lysates were subjected to Phos-tag SDS-PAGE followed by western blotting to assess the phosphorylation status of ADAR1p110. A reduction in the slower-migrating phosphorylated form of ADAR1p110 was observed upon CDK13 knockdown, confirming its role in mitotic phosphorylation.

To determine whether ADAR1p110 is post-translationally modified during mitosis, we performed phosphopeptide mapping by mass spectrometry using 3×Flag-tagged ADAR1p110 purified from 293T cells under either Async or Msync conditions **(Fig. 6B)**. Detailed mass spectrometry spectra and peptide coverage are shown in **Supplemental Figure 3**. Comparative analysis of peptide coverage revealed several phosphorylation-enriched regions that were strongly detected under Msync conditions but absent or diminished in asynchronous samples. These phosphorylation sites were clustered within a conserved serine/threonine-rich domain, notably encompassing serine 614. This site is of particular interest as it conforms to a CDK13 consensus motif (Fan et al. 2020).

To evaluate the functional significance of ADAR1p110 phosphorylation during mitosis, we performed rescue experiments in HeLa cells in which endogenous ADAR1 was depleted using siRNA targeting the 3′-UTR **(Fig. 6C)**. Cells were then transfected with mCherry-tagged ADAR1p110 constructs, including WT, phospho-mimetic mutants (3×D: three serine/threonine sites substituted with aspartic acid; S614D: serine 614 substituted individually), and phospho-deficient mutants (3×A and S614A). Phospho-histone H3 (Ser10) levels were analyzed by western blotting as a readout of mitotic accumulation. As expected, ADAR1 knockdown led to increased levels of phospho-histone H3 (S10), indicating mitotic arrest. Expression of phospho-mimetic constructs (3×D and S614D) substantially reduced phospho-H3 (S10) levels, consistent with restoration of normal mitotic progression. By contrast, neither WT nor phospho-deficient forms (3×A and S614A) alleviated the accumulation of mitotic markers. In addition to their effects on mitotic progression, these phospho-mutants exhibited distinct catalytic activities. Inosine deamination assays using recombinant ADAR1p110 variants showed that the phospho-mimetic form (3×D) had higher editing efficiency than WT, while the phospho-deficient mutant (3×A) exhibited impaired enzyme activity **(Supplemental Fig.4)**. These results demonstrate that ADAR1p110 undergoes mitosis-specific phosphorylation, particularly at serine 614, which is required for its function during mitotic progression. The results of phospho-specific mobility shift assays and phosphatase treatment, and the functional rescue of mitotic arrest following ADAR1p110 depletion by the S614D mutant collectively support a regulatory mechanism in which phosphorylation at S614 acts as a critical switch for ADAR1p110 activation in M phase.

To identify the kinase(s) responsible for phosphorylation of ADAR1p110 during mitosis, we treated HeLa cells with selective inhibitors targeting PLK1 (BI2536 and GSK461364) or CDK12/13 (SR-4835 and THZ531; **Fig. 6D)**. Cells were analyzed under both Async and Msync conditions using Phos-tag SDS-PAGE to resolve phosphorylated ADAR1p110 species. Treatment with the PLK1 inhibitors BI2536 and GSK461364 resulted in a partial reduction in mobility-shifted phosphorylated bands of ADAR1p110, suggesting that PLK1 contributes to—but is not solely responsible for—its mitotic phosphorylation. By contrast, CDK12/13 inhibitors produced a much more pronounced effect. Both SR-4835 and THZ531 led to a dose-dependent loss of phosphorylated ADAR1p110 bands, nearly abolishing the shift at higher concentrations.

These findings indicate that CDK13 and/or CDK12 are the principal kinases mediating ADAR1p110 phosphorylation during mitosis, with PLK1 playing a supporting or secondary role. To directly assess the role of CDK13 in mediating ADAR1p110 phosphorylation during mitosis, we performed siRNA knockdown of CDK13 in HeLa cells **(Fig. 6E)**. Cells were synchronized in mitosis using nocodazole and transfected with either CDK13-specific siRNA (siCDK13) or a non-targeting control (siNC1). Phos-tag SDS-PAGE followed by immunoblotting revealed a substantially lower level of phosphorylated ADAR1p110 in cells treated with siCDK13 than in mitotically synchronized controls. The diminished mobility-shifted bands indicate that phosphorylation of ADAR1p110 is strongly dependent on CDK13 activity. These results reinforce our pharmacological findings and provide genetic evidence that CDK13 is a key upstream kinase responsible for ADAR1p110 phosphorylation during mitosis. Notably, computational kinase prediction using PhosphoSitePlus™ (Cell Signaling Technology) also identifies CDK13 as the top candidate kinase for ADAR1 S614 phosphorylation with 95% confidence, further supporting our experimental findings.

## Discussion

ADAR1 is best known as a key regulator of A-to-I RNA editing, primarily acting in the cytoplasm to prevent aberrant innate immune responses by targeting dsRNAs. However, this study revealed a distinct nuclear role for the ADAR1p110 isoform during mitosis—separate from its canonical editing function. Using a combination of live cell imaging, genomic profiling, proteomics, and biochemical approaches, we demonstrate that ADAR1p110 dynamically localizes to mitotic chromatin, interacts with cohesin components such as SMC3, and undergoes mitosis-specific phosphorylation that is essential for its stability and functional engagement during cell division.

Loss of ADAR1 leads to profound mitotic phenotypes, including metaphase arrest, activation of DNA damage markers (γH2AX, phospho-RPA32, phospho-DNA-PKcs), and induction of apoptotic signaling (cleaved PARP). These defects are accompanied by accumulation of mitotic regulators such as phospho-Histone H3 (Ser10), Cyclin B1, phospho-PLK1, and phospho-Aurora kinases. Collectively, these results suggest that ADAR1p110 is required for timely mitotic progression and genome stability.

Importantly, the results also revealed that ADAR1p110 is enriched at centromeric α-satellite DNA during mitosis. Genome-wide mapping using DIP-seq and targeted validation by DIP-qPCR both confirmed mitosis-specific recruitment of ADAR1p110 in centromeric regions, particularly those enriched in R-loops. These RNA:DNA hybrid structures are increasingly recognized as functional regulators of centromeric architecture, and their formation was further enhanced under conditions of mitotic synchronization and ADAR1 overexpression. Given the known affinity of ADAR1 for RNA:DNA hybrids, this suggests that ADAR1p110 is selectively recruited to centromeric R-loop-containing regions during mitosis.

Importantly, ADAR1p110 also interacts with SMC3, a core cohesin subunit, suggesting that it may contribute to maintaining centromeric structure or sister chromatid cohesion. The cohesin complex is essential for mitotic fidelity, and recent reports indicate that R-loops help anchor cohesin at specific genomic loci (Zhang et al. 2023; Porter et al. 2023). Dual interaction of ADAR1p110 with centromeric R-loops and SMC3 places it at the nexus of these regulatory modules. We speculate that ADAR1p110 may reinforce cohesin retention or function at centromeres, particularly during metaphase, thereby contributing to faithful chromosome segregation.

Mechanistically, we identified phosphorylation of ADAR1p110 at serine 614 (S614) as a key mitosis-specific regulatory event. Phos-tag electrophoresis and mass spectrometry confirmed robust phosphorylation of ADAR1p110 at S614 specifically during mitosis. This modification is essential for protein stability because the S614A phospho-deficient mutant was unstable during mitosis and rescued by proteasome inhibition. By contrast, the phospho-mimetic S614D mutant restored mitotic progression in ADAR1-depleted cells, highlighting the functional necessity of this post-translational modification.

Through kinase inhibition and RNA interference (RNAi) experiments, we identified CDK13 as the principal kinase responsible for S614 phosphorylation. Inhibition of CDK13 using SR-4835 or THZ531 abolished the mobility shift of ADAR1p110 observed on Phos-tag gels, and CDK13 knockdown produced similar effects. While PLK1 inhibitors partially reduced phosphorylation, CDK13 appears to be the predominant kinase responsible for this phosphorylation.

Interestingly, CDK13 is not only an upstream kinase of ADAR1, but also a well-established RNA editing target of ADAR1 (Ramírez-Moya et al. 2021). Multiple editing sites in the CDK13 transcript, including Q103R and Q113R, result in altered protein-coding potential, and have been shown to promote nucleolar localization and cellular proliferation in cancer models. This bidirectional relationship—in which CDK13 edits and is edited by ADAR1—highlights a unique post-transcriptional and post-translational feedback axis in mitotic control.

*In vitro* RNA editing assays revealed that the phospho-mimetic 3×D mutant of ADAR1p110 exhibited accelerated early-phase A-to-I activity compared with WT and phospho-deficient versions. Although all variants ultimately reached similar editing saturation levels, this suggests that phosphorylation may enhance the initial catalytic efficiency of ADAR1p110. While not central to its mitotic chromatin function, this effect may be relevant in environments where rapid RNA processing is required, such as during mitotic reentry or transcriptional shutdown.

Together, these findings position ADAR1p110 as a mitosis-specific chromatin-associated factor that links R-loop recognition to cohesin-dependent chromosome cohesion. Phosphorylation at S614 by CDK13 activates ADAR1p110 for chromatin binding and protects it from proteasomal degradation, enabling it to stabilize centromeric architecture during chromosome segregation. This work expands the known roles of ADAR1 beyond RNA editing and establishes a regulatory network involving CDK13, ADAR1p110, and centromeric chromatin that supports mitotic fidelity and genome integrity.

## Methods

### Cell culture

HeLa human ovarian carcinoma (ATCC CCL-2), HEK293T human embryonic kidney (ATCC CRL-11268), and HCT116 human colon carcinoma (ATCC CCL-247) cells were cultured in Dulbecco’s modified Eagle’s medium (DMEM; Nacalai) supplemented with 10% fetal bovine serum (FBS; Gibco) and 1% penicillin-streptomycin (FUJIFILM Wako), and maintained at 37°C in a humidified incubator with 5% CO₂. These cell lines were free of mycoplasma contamination.

For inducible overexpression of ADAR1p110, a previously established HeLa Tet-On stable cell line expressing 1×FLAG-ADAR1p110 was used. Doxycycline (Clontech) was added to the culture medium at a final concentration of 2.0 μg/mL for 4 days to induce protein expression.

### siRNA transfection for ADAR1 knockdown

siRNAs targeting ADAR1 were transfected into cells using Lipofectamine RNAiMAX (Thermo Fisher Scientific) according to the manufacturer’s protocol. Cells were seeded 1 day prior to transfection at a density of 4.0 × 10⁴ cells/mL. On the day of transfection (Day 0), siRNAs were transfected at a final concentration of 2 nM using the following complex preparation: siRNA (10 mM, 2.4 μL) was diluted in 1 mL of Opti-MEM (Thermo Fisher Scientific) to form solution A; 20 μL of RNAiMAX was diluted in 1 mL of Opti-MEM to form solution B. The two solutions were mixed and incubated for 5 min at room temperature before being added to cells in 10 cm dishes (final culture volume 10 mL). Cells were harvested 72 h after transfection (Day 3).

The following siRNAs were used:

Negative control siRNA (siNC1; AM4611, Thermo Fisher Scientific)

siADAR1-1 (53004, Thermo Fisher Scientific, custom):

Sense: 5′-AGAGGUAGGUCGUAGCAUUtt-3′
Antisense: 5′-AAUGCUACGACCUACCUCUct-3′

siADAR1-2 (53005, Thermo Fisher Scientific, custom):

Sense: 5′-GAUGGAUUAGGGUGUGUCAtt-3′
Antisense: 5′-UGACACACCCUAAUCCAUCtg-3′

### Flow cytometric analysis of mitotic cells

HeLa cells were transfected with either control siRNA or ADAR1-targeting siRNA as described above. At 48, 60, and 72 h post-transfection, cells were harvested using enzyme-free cell dissociation buffer (Thermo Fisher Scientific), washed once with cold phosphate-buffered saline (PBS), and fixed in ice-cold 100% methanol for 15 min on ice. After fixation, cells were washed three times with PBS and stained for 1 h at room temperature in PBS containing Alexa Fluor 488-conjugated anti-phospho-histone H3 (Ser10) antibody (CST, #3465S, 1:50 dilution), then stained with 5 μM DRAQ5 (CST, #4084). Samples were analyzed on a BD LSR II flow cytometer (BD Biosciences), and data were processed using FlowJo software (FlowJo, LLC).

### Cell cycle synchronization and inhibitor treatment

For mitotic synchronization (Msync), HeLa cells were treated with nocodazole (Sigma) at a final concentration of 0.1 μg/mL for 16 h. After treatment, mitotic cells were collected by harvesting the rounded, detached cells from the culture medium. To arrest cells in S phase, thymidine (Fujifilm Wako, 207-19421) was added at a final concentration of 2.5 mM for 24 h. After treatment, cells were washed three times with PBS and released into fresh medium. Cells were harvested 10 h after release.

For inhibitor-based experiments, cells were synchronized using a two-step protocol. Thymidine (2.5 mM) was first added for 24 h, followed by three PBS washes and a 2 h release into fresh medium. Nocodazole (0.1 μg/mL) was then added together with kinase inhibitors, including BI-2536 (PLK1 inhibitor, Selleck), GSK461364 (PLK1 inhibitor, MedChemExpress), SR-4835 (CDK12/13 inhibitor, Selleck), or THZ531 (CDK12/13 inhibitor, Selleck). Cells were incubated with nocodazole and inhibitors for 18 h before harvesting.

### Antibodies

The following primary antibodies were used for western blotting and IP: anti-ADAR1 (Santa Cruz, sc-73408, 1:1000), anti-GAPDH (Cell Signaling Technology [CST], #5174, 1:4000), anti-γH2AX (S139) (CST, #9718, 1:500), anti-phospho-Histone H3 (S10) (CST, #53348, 1:1000), anti-phospho-Aurora A/B/C (Thr288/232/198) (CST, #14475, 1:1000), anti-Aurora A (CST, #2914, 1:1000), anti-Aurora B (Abcam, ab2254, 1:1000), anti-phospho-PLK1 (T210) (CST, #9062, 1:1000), anti-PLK1 (Santa Cruz, sc-17783, 1:500), anti-CDC25C (CST, #4688, 1:1000), anti-α-tubulin (CST, #2125, 1:1000), anti-SMC1 (CST, #6892, 1:1000), anti-SMC2 (CST, #5392, 1:1000), anti-SMC3 (CST, #5696, 1:1000), anti-RAD21 (CST, #4321, 1:1000), and anti-DHX9 (Abcam, ab26271, 1:1000). Secondary antibodies were anti-mouse IgG (Jackson ImmunoResearch, 715-035-151, 1:3000) and anti-rabbit IgG (Jackson ImmunoResearch, 711-035-152, 1:3000).

### Western blotting

Cells were lysed in 1.5× Laemmli sample buffer supplemented with protease inhibitors (cOmplete ultra/mini EDTA-free, Roche), phosphatase inhibitors (PhosSTOP, Roche), and 100× Benzonase (final: 0.5 U/μL Benzonase, 1 mM MgCl₂), followed by incubation at room temperature for 5 min. Lysates were then heat-denatured at 98°C for 3 min (or 10 min for crosslinked samples). Proteins were separated on 4−20% Mini-PROTEAN TGX precast gels (Bio-Rad) by SDS-PAGE and transferred to polyvinylidene fluoride (PVDF) membranes (Merck) at 100 V for 1 h. Membranes were blocked with 1% bovine serum albumin (BSA) in PBST (0.05% Tween-20) for 30 min at room temperature. Primary antibodies were diluted in 1% BSA-PBST and incubated with membranes for 2 h at room temperature. Secondary antibody incubation was performed using the Snap i.d. 2.0 system (Millipore) for 10 min at room temperature. Protein signals were detected using Chemi Lumi One L reagent (Nacalai) and imaged using an iBright imaging system (Thermo Fisher Scientific).

### Immunoprecipitation

For immunoprecipitation (IP), Anti-FLAG M2 magnetic beads (Sigma, M8823) were used. Beads were prewashed three times by gentle inversion in TBS buffer comprising 50 mM Tris-HCl (pH 7.4), 150 mM NaCl, 0.1% NP-40, and 5% BSA, then incubated overnight at 4°C on a rotator in the same buffer. After incubation, beads were washed an additional three times before use.

Cells used for IP were crosslinked with 0.3% formaldehyde in PBS for 10 min at room temperature with gentle mixing. The crosslinking reaction was quenched by adding 125 mM glycine. Cells were washed with PBS, pelleted by centrifugation, and lysed in RIPA+e buffer comprising 50 mM Tris-HCl pH 7.4, 150 mM NaCl, 1 mM EDTA, 1% NP-40, 0.5% sodium deoxycholate, 0.1% SDS supplemented with protease inhibitor cocktail (Roche), PhosSTOP (Roche), and RNasin Plus RNase Inhibitor (Promega) at a concentration of 1×10⁷ cells/mL. Lysates were incubated at 4°C for 20 min with gentle inversion, followed by sonication using a probe-type sonicator (SONICS) with a 3 mm tip. Sonication was performed at 60% amplitude with 4 s on / 45 s off cycles, repeated 25 times. Lysates were then cleared by centrifugation at 15,000 × g for 15 min at 4 °C and cleared lysates were incubated with prewashed FLAG magnetic beads overnight at 4°C with rotation. After incubation, beads were washed three times with RIPA+e buffer by gentle inversion, followed by a 15 min rotation in RIPA+e buffer containing protease and phosphatase inhibitors. A final set of three washes was performed using NP-40+e buffer (50 mM Tris-HCl pH 7.4, 150 mM NaCl, 1 mM EDTA, 1% NP-40). Beads were resuspended in TBS buffer and stored at −30°C until further processing.

### DNA purification following IP

Beads were washed with TE buffer (pH 8.0) and incubated with elution buffer (1 M Tris-HCl pH 8.0, 0.5 M EDTA, 10% SDS) at 65°C for 30 min with shaking at 1400 rpm. Eluates were transferred to new tubes and treated with NaCl to a final concentration of 0.2 M. Samples were incubated at 65°C for 15 h for reverse crosslinking. Next, 50 mM Tris-HCl (pH 8.0) was added to a final concentration of 25 mM, along with RNase I (Invitrogen) at a total amount of 100 units. The mixture was incubated at 37°C for 2 h. Proteinase K (final: 0.25 mg/mL) was then added and incubated at 55°C for 2 h. DNA was purified using a MinElute PCR Purification Kit (QIAGEN) according to the manufacturer’s instructions.

### Plasmid construction

The fluorescent fusion constructs mCherry-ADAR1p110 and EYFP-ADAR1p110 were generated as described previously (Sakurai et al. 2017). Constructs containing alanine or aspartic acid mutations at four conserved phosphorylation sites (S599, T601, S605, and S614) were generated by PCR-based site-directed mutagenesis using mutation-specific primers **(Supplemental Table 1).**

The EGFP-α-Tubulin plasmid (pIRESneo-EGFP-α-Tubulin) was obtained from Addgene. The dsRNA-binding-defective mutant (mCherry-ADAR1p110-EAA) and the deaminase-deficient mutant (EYFP-ADAR1p110-E912A) were also generated by PCR-based mutagenesis as previously reported (Sakurai et al. 2017).

### Fluorescence microscopy

Transfection of fluorescent protein-tagged plasmids into HeLa cells was carried out using Lipofectamine 3000 (Thermo Fisher Scientific) as previously described (Sakurai et al. 2017). Cells were seeded into μ-Dish 35-mm high ibiTreat dishes (Ibidi), transfected with the respective plasmids, and incubated at 37°C for 24−36 h.

Cells were fixed in 4% formaldehyde in PBS containing 1 μg/mL Hoechst 33342 (Thermo Fisher Scientific) for 10 min at room temperature. Residual formaldehyde was quenched by two washes with 0.1 M glycine in PBS, followed by two additional PBS washes. Fluorescence images were acquired using a Leica TCS SP5 DMI6000 CS confocal microscope, equipped as previously described (Sakurai et al. 2017). Images were analyzed using Leica LAS AF software and ImageJ (NIH).

### DRIP-seq sample preparation

Genomic DNA was extracted using a Blood & Cell Culture DNA Maxi Kit (QIAGEN) and resuspended at 50 μg in 10 mM Tris-HCl (pH 8.5) and 300 mM NaCl. DNA was fragmented into 200−500 bp pieces by sonication using a probe-type sonicator (SONICS) with a 2 mm tip, applying 4 s on / 45 s off cycles at 40% amplitude. To perform DNA:RNA hybrid IP, sheared DNA (210 μL) was mixed with an equal volume of 2× DRIP dilution buffer containing 100 mM Tris-HCl (pH 7.4), 10 mM EDTA, 2.0% NP-40, and 0.2% sodium deoxycholate. S9.6 antibody (20 μg) prebound to 100 μL of Protein A magnetic beads (Thermo Fisher Scientific) was then added to the DNA mixture. Samples were incubated on a rotator at 4°C for 2 h to overnight. All buffers were prechilled and all steps were performed at 4°C. After IP, magnetic beads were collected using a magnetic stand and sequentially washed at 4°C. Specifically, beads were first washed twice with 1 mL of low-salt DRIP buffer containing 50 mM Tris-HCl (pH 7.4), 150 mM NaCl, 5 mM EDTA, 1.0% NP-40, and 0.1% sodium deoxycholate. This was followed by two washes with high-salt DRIP buffer containing 50 mM Tris-HCl (pH 7.4), 500 mM NaCl, 5 mM EDTA, 1.0% NP-40, and 0.1% sodium deoxycholate, and then two washes with DRIP Li buffer containing 50 mM Tris-HCl (pH 8.0), 250 mM LiCl, 1 mM EDTA, 0.5% NP-40, and 0.5% sodium deoxycholate. For the first three washing steps, beads were removed from the magnet and fully resuspended before being recollected. Subsequent washes were performed on the magnetic stand, once with TE + salt buffer (100 mM Tris-HCl, pH 8.0; 10 mM EDTA; 50 mM NaCl), and once with TE buffer (100 mM Tris-HCl, pH 8.0; 10 mM EDTA), with gentle inversion. Residual buffer was removed by centrifugation at 960 × g for s. Beads were eluted in 200 μL of freshly pre-warmed elution buffer containing 50 mM Tris-HCl (pH 8.0), 10 mM EDTA, and 1.0% SDS, followed by incubation at 65°C for 30 min with shaking at 1,400 rpm. Samples were briefly centrifuged and placed on a magnetic stand to recover the eluate. To eliminate potential bead contamination, the eluate was transferred to a new tube and centrifuged at 16,000 × g for 1 min at room temperature. To remove proteins and stabilize RNA, 4 μL of Proteinase K (20 mg/mL) and 4 μL of RNasin Plus RNase Inhibitor (Promega) were added to each eluate, followed by incubation at 42°C for 30 min. DNA:RNA hybrids were purified using a MinElute PCR Purification Kit (QIAGEN) according to the manufacturer’s instructions. To ensure optimal pH during column binding, 10 μL of 3 M sodium acetate (pH 5.2) was added before the binding step as recommended. DNA:RNA complexes were eluted in 12 μL of 5 mM Tris-HCl (pH 8.5).

### Next-generation sequencing (NGS) and bioinformatics analysis

Library preparation and NGS using an Illumina NovaSeq 6000 platform were outsourced to Azenta Life Sciences.

DIP-seq data were processed to remove adapters using Trim_Galore (version 0.6.10; https://www.bioinformatics.babraham.ac.uk/projects/trim_galore/) with parameters set to -q 20 --illumina --length 20 --paired --trim-n (Krueger et al. 2023). Subsequently, trimmed reads were aligned to the T2T genome (CHM13v2.0) (Rhie et al. 2023) using BWA-MEM (version 0.7.17-r1188) (Li 2013) with the settings -h 100 -q. Note that only one hit for a multi-mapping read is recorded in the resulting BAM files. The BAM files were then sorted and indexed using Samtools (version 1.6) (Li et al. 2009). Duplicate reads within BAM files were identified and removed using Picard (version 2.20.4) (Broad 2019). Peaks were extracted from the deduplicated BAM files using MACS2 peakcall (version 2.2.6; https://github.com/macs3-project/MACS) with the parameters -f BAMPE -g hs (Feng et al. 2012). Finally, we used the RIdeogram library (https://github.com/TickingClock1992/RIdeogram) to visualize the distribution of peaks on idiograms (Hao et al. 2020). Bedtools intersect (Quinlan and Hall 2010) was employed to calculate the length of overlap between peaks and repeat regions (RepeatMasker v4.1.2p1.2022Apr14 (Smit et al. 2015)), from which we computed the density of each repeat class within the peak regions. For sequences corresponding to peaks in each group, we extracted enriched sequence motifs using the parameters -dna -nostatus -time 14400 -mod anr -nmotifs 10 -minw 6 -maxw 12 -objfun classic -revcomp -markov_order 0 in the MEME suite (version 5.0.5) (Bailey et al. 2015).

For the DRIP-seq data, we aligned the reads to the genome using Bowtie2 with the parameters --end-to-end --very-sensitive --no-mixed --no-discordant -I 10 -X 700 to generate SAM files (Langmead and Salzberg 2012). After sorting and removing duplicates from the SAM files with Picard, we converted them to BAM format using Samtools for downstream analysis. MACS2 peakcall was used with the parameters -f BAMPE --broad --broad-cutoff 0.1 to extract R-loop peaks from the BAM files. Bedtools intersect (Quinlan and Hall 2010; https://bedtools.readthedocs.io/) was then applied to quantify the number of reads within these peaks and to identify those overlapping with centromeric regions.

### qPCR analysis of immunoprecipitated DNA

To assess the enrichment of ADAR1p110 at centromeric regions, qPCR was performed using DNA obtained from anti-FLAG IP experiments. DNA was purified as described above and used directly for qPCR analysis. qPCR was carried out using Luna Universal qPCR Master Mix (New England Biolabs) and gene-specific primers targeting centromeric sequences of chromosomes 1, 5, and 19. Reactions were performed on a QuantStudio 3 Real-Time PCR System (Applied Biosystems) according to the manufacturer’s instructions. Relative enrichment was calculated based on the percentage of input DNA, and signals were compared across negative control (no FLAG-ADAR1), asynchronous FLAG-ADAR1 expression, and mitotically synchronized FLAG-ADAR1 expression conditions. The following primers were used:

Chromosome 1 centromere: forward 5′-GCCGCTTTGAGGTCAATGGTA-3′, reverse 5′-CACTTGCAGACTTTACAAACAGAGT-3′

Chromosome 5 centromere (Slee et al. 2012) : forward 5′-TAGACAGAAATATTCTCACAATCGT-3′, reverse 5′-GCCCTCAAAGCGCTCCAAG-3′

Chromosome 19 centromere: forward 5′-GTGGATATTCAGACCTCCTT-3′, reverse 5′-TGTTTCAAGTCTGCTCTGTGTA-3′

### Liquid chromatography tandem mass spectrometry (LC-MS/MS) analysis of phosphorylated ADAR1p110

HEK293T cells (ATCC CRL-11268) stably expressing FLAG-tagged ADAR1p110 were synchronized in mitosis using nocodazole. FLAG-ADAR1p110 proteins were immunoprecipitated and processed for phosphoproteomic analysis by LC-MS/MS as previously described (Sakurai et al. 2017).

### Phos-tag SDS-PAGE

Phosphorylated proteins were separated using Phos-tag SDS-PAGE with Mn²⁺-dependent phosphate affinity. Prior to gel loading, 10 mM MnCl₂ was added to the sample solution to a final concentration of 2 mM. The separating gel (6.75% acrylamide) was prepared by incorporating 5 μM Phos-tag acrylamide (Wako, AAL-107; final concentration 2.5 μM) and 10 mM MnCl₂ (final concentration 0.125 mM). After loading, electrophoresis was performed using SDS running buffer at 100 V through the stacking gel, followed by separation at 60 mA through the resolving gel. To avoid reduced protein transfer efficiency due to Mn²⁺, the gel was washed sequentially after electrophoresis for 10 min in SDS running buffer containing 2.5 mM EDTA, 10 min in transfer buffer containing 2.5 mM EDTA, and 5 min in transfer buffer without EDTA. Proteins were transferred to PVDF membranes at 105 V for 1 h. Subsequent immunodetection was performed as described above.

## Data access

### Supplementary information

**Supplemental_Fig._S1.pdf**

**Supplemental_Table_S1.xls**

## Acknowledgments

This work was supported by JSPS KAKENHI [JP18H06054, JP19H03159, JP22H02548, JP23K23812, JP22K15093, JP24K21326, JP23H00509]; AMED [JP25gm7010005, JP25ae0121055, JP25ama121055]; Takeda Science Foundation; Astellas Foundation for Research on Metabolic Disorders; The Sumitomo Foundation (210858); The Uehara Memorial Foundation; Senri Life Science Foundation; Japan Cancer Society; The Yasuda Medical Foundation; The Ichiro Kanehara Foundation for the Promotion of Medical Sciences and Medical Care; MSD Life Science Foundation, Public Interest Incorporated Foundation; Mochida Memorial Foundation for Medical and Pharmaceutical Research.

## Author contributions

M.S. and M.K. designed the research. Y.Y., M.K., K.H., and M.S. performed most cellular and biochemical experiments. Some early-stage experiments were conducted under the guidance of K.N. C.Z. performed sequencing data processing under supervision of M.H. Y.Y. and M.S. wrote the manuscript with input from all authors. M.S. supervised the overall project.

